# BioXNet: a biologically inspired neural network for deciphering anti-cancer drug response in precision medicine

**DOI:** 10.1101/2024.01.29.576766

**Authors:** Jiannan Yang, William Ka Kei Wu, Rina Yee Man Hui, Ian Chi Kei Wong, Qingpeng Zhang

**Affiliations:** School of Public Health, LKS Faculty of Medicine, The University of Hong Kong, Hong Kong SAR, China; The Laboratory of Data Discovery for Health, Hong Kong SAR, China; Department of Anaesthesia and Intensive Care, Chinese University of Hong Kong, Hong Kong SAR, China; Centre of Cancer Medicine, School of Clinical Medicine, LKS Faculty of Medicine, The University of Hong Kong, Hong Kong SAR, China; Department of Pharmacology and Pharmacy, LKS Faculty of Medicine, The University of Hong Kong, Hong Kong SAR, China; Musketeers Foundation Institute of Data Science, The University of Hong Kong, Hong Kong SAR, China

## Abstract

Accurate prediction of anti-cancer drug responses in preclinical and clinical studies is crucial for drug discovery and personalized medicine. While machine learning models have demonstrated promising prediction accuracy in this task, their translational value in cancer therapy is constrained by the lack of model interpretability and insufficient patients’ data with genomic profiles to calibrate models. The rich cell line data has the potential to supplement patients’ data, but the difference between the drug response mechanisms in cell lines and human body needs to be characterized quantitatively. To address these challenges, we proposed the BioXNet, which captures drug response mechanisms by seamlessly integrating drug target information with genomic profiles (genetic and epigenetic modifications) into a single biologically inspired neural network. BioXNet exhibited superior performance in drug response prediction tasks in both preclinical and clinical settings. An analysis of BioXNet’s interpretability revealed its ability to identify significant differences in drug response mechanisms between cell lines and the human body. Notably, the key factor of drug response is the drug targeting genes in cell lines but methylation modifications in the human body. Furthermore, we developed an online human-readable interface of BioXNet for drug response exploration by medical professionals and laymen. BioXNet represents a step further towards unifying drug, cell line and patients’ data under a holistic interpretable machine learning framework for precision medicine in cancer therapy.

## Introduction

The treatment of cancer has long been a challenging endeavor. Recent development of personalized cancer therapies^1–3^ has become a focal point in oncology research, aiming to improve patient outcomes by tailoring treatment plans to individual needs. A crucial aspect of this endeavor is the accurate prediction of drug response, which serves as a foundation for selecting effective and appropriate therapeutic options for cancer patients. Predicting drug response with high precision enables clinicians to maximize the chance of successful treatment, thereby enhancing patient care and overall survival rates.

There has been a substantial effort in introducing machine learning approaches^4^ to accurately predict drug response. Existing research has employed machine learning models such as ElasticNet^5^ and random forest^6,7^ to predict drug response in preclinical settings. These preclinical settings typically involve human cancer cell lines, which are derived from patient tumor issues and cultured *in vitro* and play a fundamental role in cancer therapy research^8–10^. Recent studies^11–14^ have developed various computational approaches to extend the drug response prediction task to clinical settings. While these computational approaches have demonstrated promising prediction performance, the limitation in interpretability restricts their ability to provide clinical insights for further applications. As cell lines cultured outside of human body, the lack of comprehensive biological systems^15,16^ may limit their ability to truly reflect the efficacy of drugs in clinical trials^17,18^. The extent to which these differences between preclinical and clinical settings can be identified and characterized by computational methods remains largely unknown. Addressing this knowledge gap has the potential to improve our understanding of the discrepancies between drug responses in cell lines and patients, thereby enhancing the development of more effective and personalized cancer treatments.

Recently, biologically inspired neural networks have been proposed to improve interpretability by simulating complex biological systems using hierarchical biological structures. Following the emergence of DCell^19^, which modeled eukaryotic cells to simulate cellular functions, this type of model has demonstrated great promise in cancer-related research, such as predicting cancer states^20^ and identifying synergistic drug combinations^21^. The extension of DCell, known as DrugCell^21^, combining a conventional neural network for chemical structure modeling with the DCell framework, identified interpretations that represented synergistic drug combination opportunities. However, these models primarily rely on genetic variation information (i.e., mutations and copy number variations) and neglect epigenetic modifications, such as methylation. These epigenetic modifications significantly contribute to cancer development^22^ and drug resistance^23,24^ due to the dysregulation of cellular function. Therefore, we hypothesize that incorporating drug target information into this biologically inspired neural network and considering both genetic and epigenetic modifications would enhance the accuracy of drug response predictions and facilitate the identification of drug response mechanisms, particularly the distinctions between these mechanisms in cell lines and human bodies.

In this study, we introduced a novel and **Bio**logically inspired e**X**plainable neural **Net**work, namely the **BioXNet**, to model drug response mechanisms by integrating drug target information with genomic profiles, which include both genetic and epigenetic modifications. BioXNet employed the hierarchical biological organization of the human body to establish its sparse model architecture, wherein nodes represent biological entities (genes and pathways), and sparse connections are derived from existing biological relationships. We validated BioXNet’s performance in predicting drug response in both preclinical and clinical contexts. The interpretability of BioXNet was demonstrated through a weighted hierarchical biological structure, wherein we employed a random walk approach to identify interpretable paths that elucidate the biological pathways associated with drug response. The variation in the explanations of drug response between cell lines and the human body highlights the differences in drug response mechanisms between preclinical and clinical settings.

## Results

### The design of BioXNet for drug response prediction

BioXNet was proposed to model the complex biological processes and provide biologically meaningful interpretations. Different from a fully connected neural network, BioXNet is a constrained system that reflects the hierarchical structure of biological processes in the human body (Reactome^25^, Figure 1c). BioXNet can capture the complex mechanisms of drug action within the human body, combining drug target information with genomic profiles to simulate these intricate processes. Unlike P-Net, a recent biologically inspired neural network for cancer state prediction^20^, which only considered genetic variations of genomic profiles, our approach incorporated both the genetic and epigenetic in genomic profiles, including methylation, copy number variation (CNV), and mutation (Figure 1a-c). Additionally, BioXNet incorporates the attention modules between layers to learn the functional associations cross layers.

**Figure 1.**
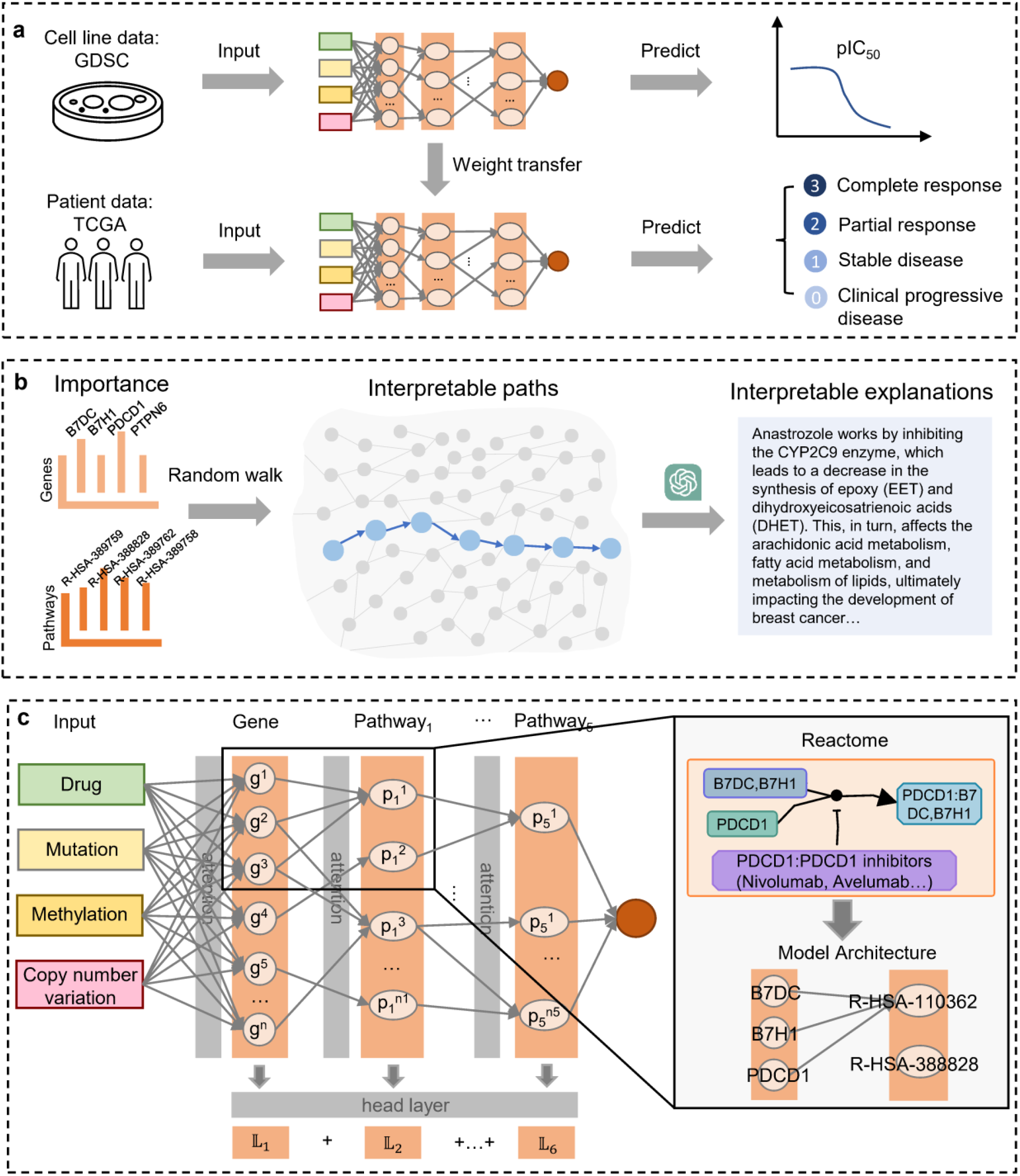
The framework of BioXNet. **a**. The application of BioXNet to preclinical (cell line data) and clinical (patient data) settings for drug response prediction. The input consists of drug target information integrated with genomic profiles. It predicts *pIC*_50_ values in preclinical datasets and Response Evaluation Criteria in Solid Tumors in clinical datasets. Weight transfer is employed by transferring trained weights from preclinical datasets to the model for clinical applications. **b**. Interpretations generated by BioXNet. DeepLIFT is adopted to compute the importance scores of nodes in each layer (genes and pathways). A random walk is adopted to identify mechanistic paths for drug response and employed a large language model (GPT-3.5 Turbo) to generate human-readable explanation of drug response. **c**. Detailed architecture of BioXNet. The neurons and neuron connections are defined by the hierarchical biological pathway network of the Reactome database. For instance, the “PD-1 binds B7DC and B7H1” pathway (R-HSA-110362) is composed of three genes (B7DC, B7H1, and PDCD1), and BioXNet contains three edges connecting these three neurons to the corresponding neuron in the subsequent layer. In contrast, the “POLB-Dependent Long Patch Base Excision Repair” pathway (R-HSA-388828) lacks any biological association with these genes, resulting in the absence of connecting edges in BioXNet.

We applied BioXNet to both preclinical and clinical settings for drug response prediction and further explored whether transferring weights could improve the performance of BioXNet. The preclinical application utilized the Genomics of Drug Sensitivity in Cancer (GDSC) cell line dataset^26^, while the clinical application employed The Cancer Genome Atlas Program (TCGA) patient dataset^27^ (Figure 1a). To enhance interpretability for medical professionals, we used DeepLIFT^28^ to compute importance scores, introduced a random walk-based method to identify interpretable paths, and employed a large language model (GPT-3.5 Turbo) to generate human-readable interpretations (Figure 1b). Further details can be found in the Methods section.

### BioXNet accurately predicts drug response in both preclinical and clinical settings

We first investigated the performance of BioXNet in drug response prediction task in the preclinical setting (GDSC dataset). As presented in Figure 2a, the performance on this task was measured using the Pearson correlation (rho) between the predicted and observed *pIC*_50_ values in 5-fold cross validation (Methods). BioXNet achieved a mean rho of 0.85 (*p* < 0.0001) for all drug-cell line pairs across all cancer types. Then, we applied BioXNet to drug response prediction in the clinical setting (TCGA dataset). As the drug response is measured using Response Evaluation Criteria in Solid Tumors (RECIST)^29^, which consists of four categories of drug response, the performance of BioXNet was measured using a micro-F1 score (Methods). BioXNet achieved a mean micro-F1 score of 0.73 (±0.16).

**Figure 2.**
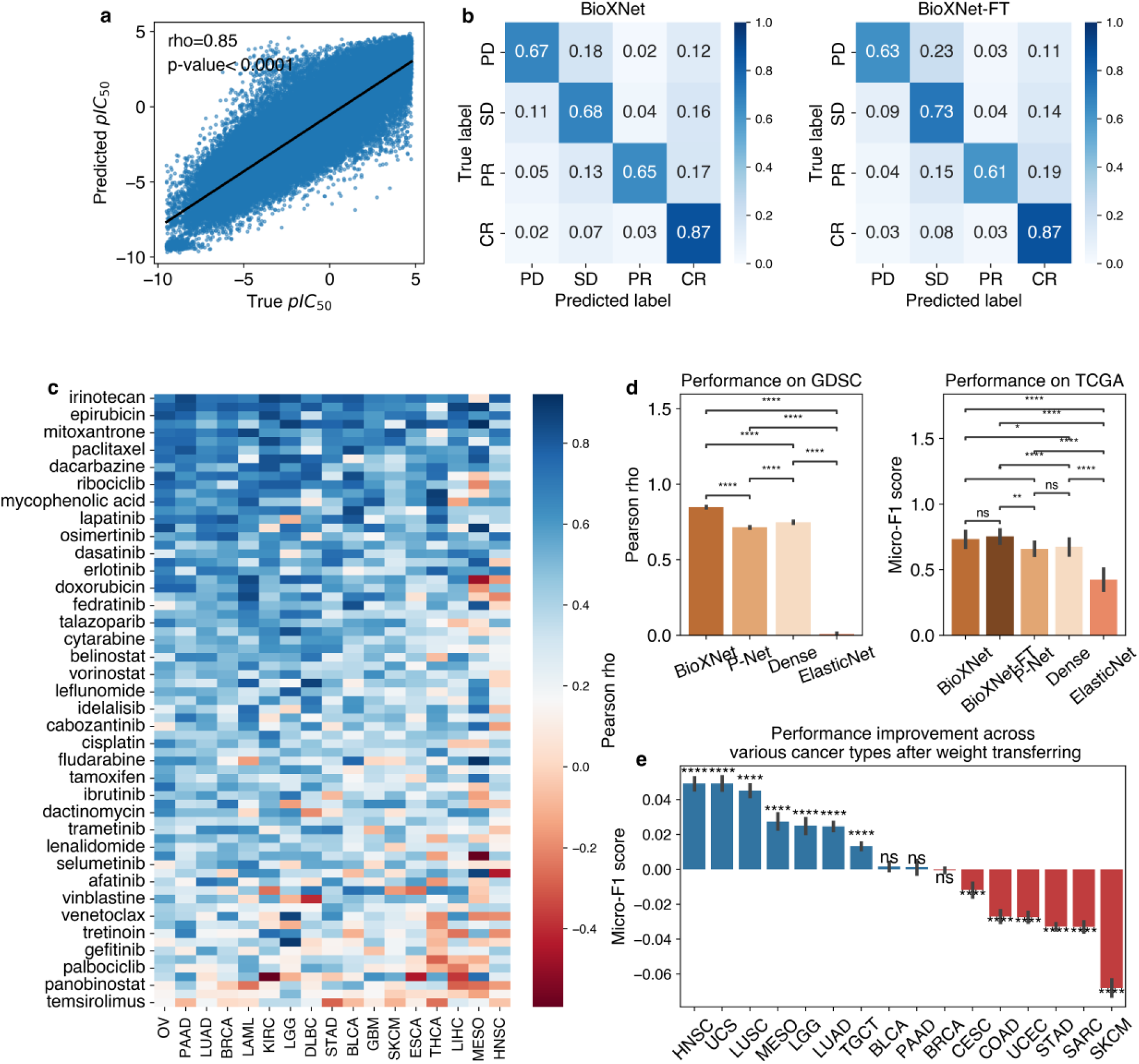
The performance of BioXNet. **a**. The Pearson’s rho between BioXNet’s prediction and the true pIC50 value in the preclinical setting (GDSC data). **b**. Confusion matrix of BioXNet and BioXNet-Finetuned in the clinical setting (TCGA data). PD, SD, PR, and CR represent drug response classifications in Response Evaluation Criteria in Solid Tumors (RECIST): Clinical Progressive Disease (PD), Stable Disease (SD), Partial Response (PR), Complete Response (CR). **c**. The heatmap of the performance of BioXNet in the preclinical setting considering different drugs and cancer types. Only FDA-approved drugs are presented. **d**. Comparison of BioXNet performance with baselines in both preclinical and clinical settings. BioXNet-FT is the abbreviation for BioXNet-FineTuned. **e**. The performance gain by weight transfer (BioXNet-FT versus BioXNet) across various cancer types in the clinical setting. The abbreviations of cancer types in **c** and **e** are OV - Ovarian serous cystadenocarcinoma; PAAD-Pancreatic adenocarcinoma; LUAD - Lung adenocarcinoma; BRCA - Breast invasive carcinoma; LAML - Acute Myeloid Leukemia; KIRC - Kidney renal clear cell carcinoma; LGG-Brain Lower Grade Glioma; DLBC - Lymphoid Neoplasm Diffuse Large B-cell Lymphoma; STAD - Stomach adenocarcinoma; BLCA - Bladder Urothelial Carcinoma; GBM - Glioblastoma multiforme; SKCM - Skin Cutaneous Melanoma; ESCA - Esophageal carcinoma; THCA - Thyroid carcinoma; LIHC - Liver hepatocellular carcinoma; MESO – Mesothelioma; HNSC - Head and Neck squamous cell carcinoma; LUSC - Lung squamous cell carcinoma; UCS - Uterine Carcinosarcoma; TGCT - Testicular Germ Cell Tumors; CESC - Cervical squamous cell carcinoma and endocervical adenocarcinoma; UCEC - Uterine Corpus Endometrial Carcinoma; COAD - Colon adenocarcinoma; SARC – Sarcoma. The *p*-values in **d** and **e** were derived from t-test with annotation legends: *ns*: *p* > 0.05; *: 0.01 < *p* ≤ 0.05; **: 0.001 < *p* ≤ 0.01; ***: 0.0001 < *p* ≤ 0.001; ****: *p* ≤ 0.0001.

BioXNet-FT (FT means fine-tuned), whose weights were transferred from BioXNet trained on the GDSC dataset, achieved a mean micro-F1 score of 0.76 (±0.14) across all cancer types in the clinical setting. Further investigation revealed that the benefits of weight transfer was inconsistent in different response types (Figure 2b), with BioXNet-FT achieving better performance on Stable Disease (SD), equal performance on Complete Response (CR), and lower performance on Progressive Disease (PD) and Partial Response (PR). This inconsistency might be due to imbalanced distributions of different cancer types and response types in the TCGA data.

We compared the performance of BioXNet with two baselines: (a) a dense model, which has the same architecture as BioXNet but uses a dense neural network with full interlayer connections. (b) P-Net^20^, a latest biologically inspired neural network for cancer state prediction. We modified the P-Net by incorporating drug target information as additional input for a fair comparison. (c) ElasticNet, a commonly used statistical model. As presented in Figure 2d, BioXNet significantly outperformed baselines on both GDSC and TCGA dataset. The performance of the classic ElasticNet was not comparable. Among the two deep learning baselines, the Dense model significantly outperformed P-Net on the GDSC dataset (median: 0.75±0.01 vs 0.71±0.01, Figure 2d) and also surpassed it on the TCGA dataset (median: 0.71±0.17 vs 0.66±0.14, Figure 2d), although the difference was not statistically significant. As P-Net utilized a similar biological-inspired architecture but without methylation information, this finding suggests that methylation information is crucial for drug response. In the case of BioXNet, we observed a similar phenomenon, wherein the performance of BioXNet without methylation also dropped (still higher than P-Net) (Figure S1a).

### The weight transfer increases the performance on specific cancer types

We found that the performance boost of BioXNet-FT compared with BioXNet was not significant across all cancer types (Figure 2d). Hence, we further investigated the performance gain by weight transfer of each cancer type with bootstrap analysis. As depicted in Figure 2e, BioXNet-FT exhibited statistically significant superior performance for specific cancer types, such as Head and Neck squamous cell carcinoma (HNSC), Lung squamous cell carcinoma (LUSC), and Uterine Carcinosarcoma (UCS), but not for other cancer types. This inconsistent finding indicates that the drug response mechanism learned from cancer cell lines are transferable to that of the human body for certain cancer types, but not others. The mapping between cell lines and the human body remains unclear, particularly concerning certain cancer types. More research on relevant genome profiles is needed to fully uncover the mechanistic differences in the drug responses in the preclinical and clinical settings.

Then, we investigated which cancer types achieved the best performance in cell lines. As shown in Figure 2c, BioXNet achieved the best performance on Ovarian serous cystadenocarcinoma (OV) (rho=0.51±0.20) and drug irinotecan (rho=0.70±0.17) on the GDSC dataset. On the TCGA dataset, BioXNet-FT achieved the best performance for Skin Cutaneous Melanoma (SKCM) and drug epirubicin (Figure S1b). We found no statistically significant correlation between the performance of BioXNet on the GDSC dataset and that of BioXNet-FT on the TCGA dataset when considering different cancer types or drugs (Figure S1c), suggesting that cancers with better performance in cell lines did not necessarily exhibit better performance in patients.

### BioXNet identifies the difference in mechanism of drug response between preclinical and clinical settings

The architecture of BioXNet allows us to investigate the contributions of different genes and biological pathways to drug response prediction. In this study, we focused on the interpretability of BioXNet on the GDSC dataset and BioXNet-FT on the TCGA dataset. We first examined the relative importance of different input data types for drug response prediction across all samples in cell lines and patients, respectively. As shown in Figure 3a, on the GDSC dataset, the input drug (median importance score: 0.025 ± 0.037) was more important for drug response prediction than the genomic profiles of cell lines. In contrast, on the TCGA dataset, methylation was identified as the primary contributor (median importance score: 0.037 ± 0.051) to patient drug response. This observation aligns with the prevalent notions that drug characteristics (e.g., targeting genes) are more decisive in drug sensitivity in cell lines, whereas the internal characteristics of patients (e.g., genomic profiles) are more influential in determining patient-specific drug response. Note that methylation was found to be the second most influential predictor of drug response in cell lines, indicating that methylation plays an important role in drug response in both cell lines and patients. Previous clinical evidence has already demonstrated that many patients had low response to certain drugs due to dysregulation of cellular function caused by epigenetic modifications^30,31^, especially the methylation of specific genes, such as CHD1^32^, MTOR^33,34^, TXK^35^, UPP1 (Figure 3g and h).

**Figure 3.**
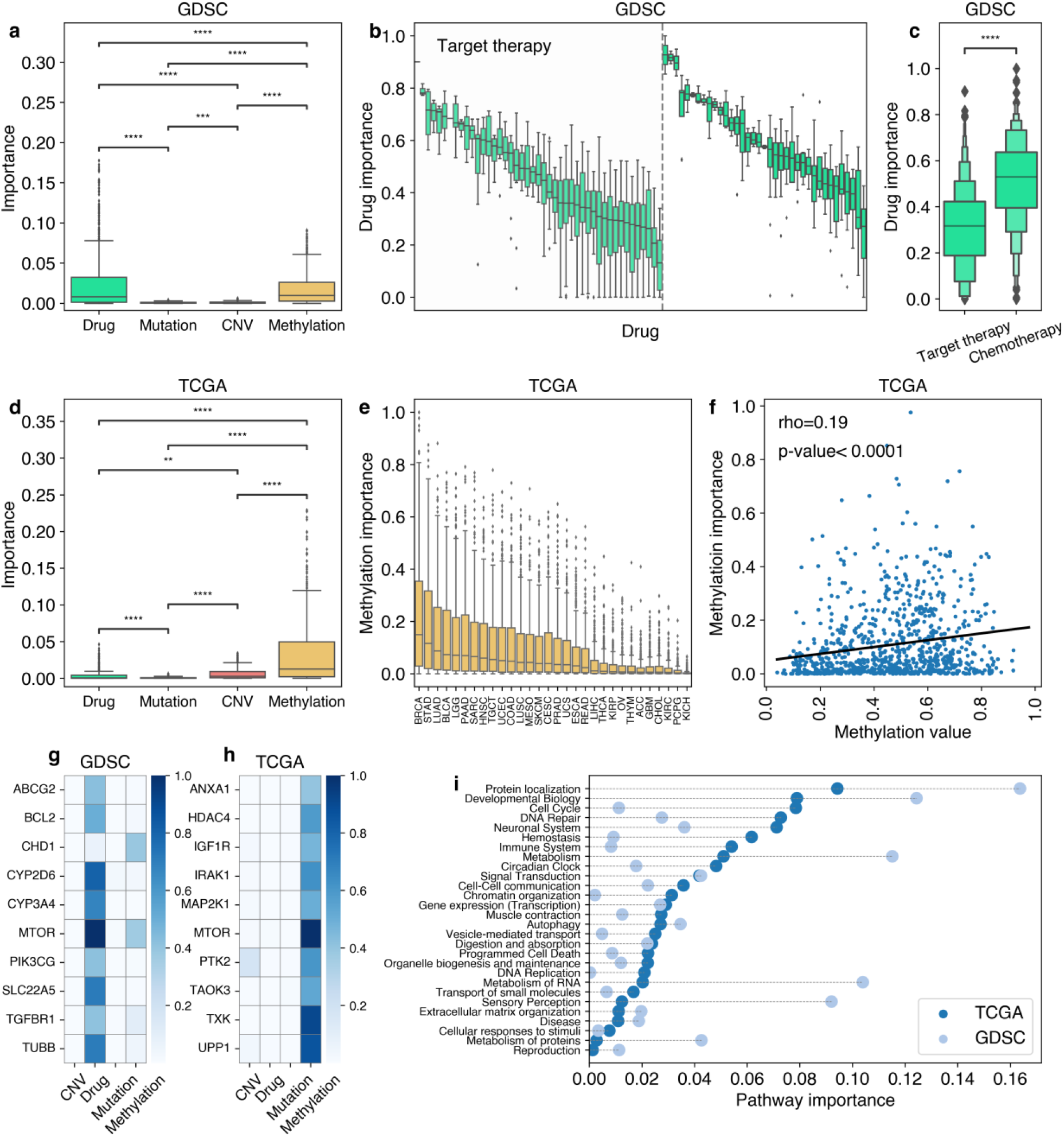
Interpretations of BioXNet in cell lines (GDSC) and BioXNet-FT in patients (TCGA). **a** and **d** are the boxplots of the importance scores of BioXNet for various input data types on the GDSC and TCGA dataset, respectively. **b**. The drug importance scores on the GDSC dataset across different drugs. **c**. The comparison of the drug importance scores for target therapy drugs and chemotherapy drugs on the TCGA dataset. **e**. The methylation importance scores on the TCGA dataset across various cancer types. **f**. The correlation of the methylation values and corresponding methylation importance scores across all cancer types for drug target genes on the TCGA dataset. **g** and **h** show the heatmaps of the importance scores of different data types across the top-10 genes on the GDSC and TCGA dataset, respectively. These genes have the top-10 importance scores when summed across four input data types. **i**. The importance scores of the highest-order biological pathways on the GDSC and TCGA dataset. The *p*-values in **a, c, d**, and **e** were derived from t-test with annotation legends: *ns*: *p* > 0.05; *: 0.01 < *p* ≤ 0.05; **: 0.001 < *p* ≤ 0.01; ***: 0.0001 < *p* ≤ 0.001; ****: *p* ≤ 0.0001.

Existing research has demonstrated that most anti-cancer drugs, particularly cytotoxic drugs, are broad-spectrum in nature^36^. Therefore, we hypothesized that chemotherapy drugs may exhibit higher drug importance scores. We categorized drugs into targeted therapies and chemotherapies based on the Food and Drug Administration (FDA) classification. On the GDSC dataset, we observed that chemotherapies attained significantly higher drug importance scores in comparison to targeted therapies. This observation was consistent with the findings on the TCGA dataset (Figure S2b-c). As expected, the presence of specific targets in the genomic profiles is crucial for determining target therapies’ drug response. As evidenced by the importance scores identified by BioXNet, the response rates of target therapies were influenced by genomic profiles (methylation in Figure 3a and 3d). Within the clinical context for each cancer type (Figure 3e), the importance of methylation was positively associated with methylation values across all genes (Figure S2a), especially for drug target genes (Figure 3f). This result suggests that, for individual patients, the methylation levels of the genes targeted by targeted therapies serves as a reliable biomarker of drug response.

Lastly, we examined the importance of biological pathways at the highest level, represented by the neurons in *Pathyway*_5_ (the last layer before the outcome layer) of BioXNet. As illustrated in Figure 3i, among the 28 such high-level biological pathways, BioXNet identified higher importance scores on the TCGA dataset in the following eight pathways: Cell Cycle, DNA Repair, Neuronal System, Hemostasis, Immune System, Chromatin Organization, Vesicle-mediated Transport, and DNA Replication. These pathways primarily involve tissue or organism-specific biological processes that are not observed in individual cell systems^15^, such as the Immune System, Neuronal System, and Vesicle-mediated Transport. Conversely, on the GDSC dataset, BioXNet identified higher importance for certain individual cell-specific biological pathways, including Protein Localization, Metabolism of RNA, and Sensory Perception.

### The interpretations of BioXNet are consistent with clinical evidence

We implemented a random walk-based approach to identify interpretable paths that effectively explain the mechanism of drug response identified by BioXNet-FT. Subsequently, we employed a Large Language Model (GPT-3.5-Turbo) to translate these paths into human readable sentences, thereby enhancing the usability of BioXNet. Please refer to our online service to explore this function (https://huggingface.co/spaces/Jayet010/bioxnet). Taking a common cancer-drug pair, breast invasive carcinoma (BRCA) and Anastrozole^37^, as an example, we examined the consistency of the interpretations derived by BioXNet with existing clinical evidence (Figure 4). Among the top 1% most frequent paths, four paths are related to the methylation of the MTOR gene (mechanistic target of rapamycin kinase). Previous research has established that the MTOR gene is closely associated with breast tumor progression^38^ and drug response rates^39^, particularly in relation to the PI3K signaling pathway^40^, which was also identified in BioXNet. Regarding the specific interpretable paths, BioXNet identified the path through MTOR methylation to CD28-dependent PI3K/Akt signaling pathway, which ultimately impacts the immune system and affects the drug response rate for breast cancer. CD28 has been recognized as a crucial pathway for immune control of breast cancer^41^. For the drug Anastrozole, BioXNet posited that its response rate is associated with its target gene CYP2C9^42^, which is involved in the synthesis of epoxy and dihydroxyeicosatrienoic acids, constituents of the arachidonic acid metabolism pathway. This pathway is connected to fatty acid metabolism and ultimately contributes to overall lipid metabolism, which is related to the progression of breast tumor^43^.

**Figure 4.**
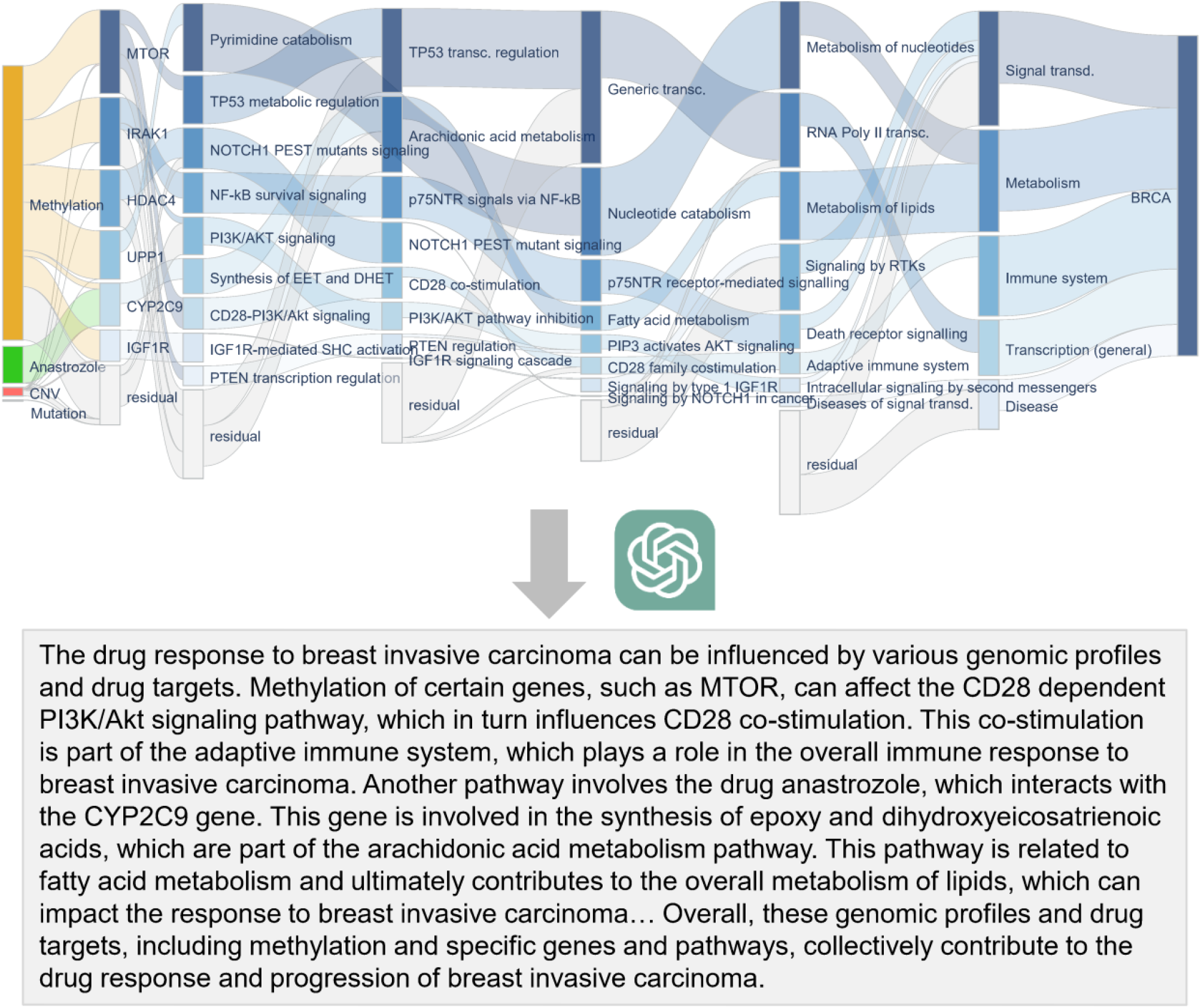
BioXNet’s interpretations of Anastrozole’s response to breast invasive carcinoma (BRCA). The Sankey plot (upper) displays the relative significance of genes and biological processes in relation to drug response identified by BioXNet-FT. The far-left and far-right layers represent input data types (drug, methylation, mutation, and copy number variation) and the outcome, respectively. Nodes in the second layer correspond to genes, while subsequent layers depict higher-order biological pathways. Node color indicates relative importance within each layer, and link width signifies the contribution from starting nodes to ending nodes, with darker colors denoting higher importance scores. The “residual” nodes in each layer refer to the set of nodes not included in the interpretable paths for that layer. We utilized GPT-3.5 Turbo to convert these paths into coherent sentences, as demonstrated in the lower figure.

## Discussion

In summary, we have developed and explored the potential of BioXNet, a biologically informed neural network that exhibits superior performance in drug response prediction across both preclinical and clinical settings. We developed several methods to enhance BioXNet’s interpretability, generating biologically meaningful interpretations that elucidate the complex mechanisms of drug response in the context of specific genomic profiles. We believe that the applications of this model can be wide-ranging: firstly, BioXNet’s direct application in predicting drug response rates for specific cancer treatments may aid patients in selecting the most effective therapies; secondly, the human-readable interpretations can help medical professionals bolster their confidence in choosing appropriate therapies; finally, the identified interpretable paths may offer insights for the development of new treatments.

A crucial aspect of BioXNet’s architecture lies not only in its biologically inspired structure but also in its distinctive design, which seamlessly integrates drug target information. Unlike DrugCell^21^, which segregated drug information from the biologically inspired neural network, BioXNet effectively captured the intricate mechanisms of drug response related to drug target genes. Moreover, BioXNet detected inconsistencies between drug responses in cell lines and patients, emphasizing that drug characteristics (targeting genes) are more deterministic for responses in cell lines, particularly for chemotherapy drugs. This observation is consistent with prior research^44,45^ indicating that most drugs exhibit uniform effectiveness or ineffectiveness across all cell lines. Additionally, we also found that weight transfer from cell line to patient dataset could only enhance performance for certain tissues. These together underscore the limitations of cell lines used in cancer therapy studies^15–18^, as their absence of comprehensive biological systems such as the immune system may constrain their capacity to accurately represent the efficacy of specific drugs in the human body. On the clinical side, epigenetic information (methylation) was identified more important for drug response prediction and the mechanism of drug response, which was also found to be highly related to drug sensitivity^46^ and resistance^30,31^ in cancer therapies. This highlights the need of incorporating methylation information for anti-cancer drug discovery and therapy design.

Although BioXNet has demonstrated promising outcomes in predicting drug responses, this *in silico* methodology necessitates further validation through *in vivo* and *in vitro* experiments to corroborate the identified critical genes and biological pathways. Moreover, BioXNet heavily depends on substantial data from drugs, cell lines or patients and previously discovered biological reactions to discern the underlying associations in cancer progression. The restricted dataset (particularly for patient data) and the incomplete drug-target information and biological reactions may result in imprecise predictions. Consequently, incorporating supplementary knowledge from patients could enhance the accuracy and dependability of BioXNet’s predictions.

## Methods

### 1. The architecture of BioXNet

BioXNet is based on a constrained neural network which is inspired by the biological processes in the human body. This architecture has already been verified effective in cancer state prediction^20^ and synergistic anti-cancer drug identification^21^. As shown in Figure 1c, the nodes in BioXNet encode some biological entities such as genes and biological pathways, and the edges within these nodes are the hierarchy biological relationships in Reactome pathway dataset. For example, the pathway “PD-1 binds B7DC and B7H1” (R-HSA-388828 in Figure 1c) is composed of three genes: B7DC, B7H1, and PDCD1, thus there are three edges starting from these three genes and targeting this pathway. For the pathway “POLB-Dependent Long Patch Base Excision Repair” (R-HSA-110862 in Figure 1c), since there is no biological relationship between these three genes and this pathway, thus different from the fully connected network, there are no edges from these three genes to this pathway in BioXNet.

To simulate the mechanism of drug action in BioXNet, given a subject (either a cell line or a patient) and a drug, the inputs to BioXNet are the genomic profiles of the subject’s genes (including methylation, mutation, and copy number variation) as well as the information about the drug’s targets. Let 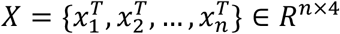 be an input array, where *n* denotes the number of genes in the first layer of BioXNet. 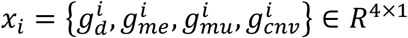 denotes the input vector of gene 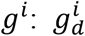 is a binary value denotes whether the input drug targets gene 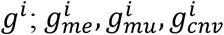 denote the methylation, mutation, and copy number variation (CNV) values for *g*^*i*^, respectively. In the first layer, we first computed a weighted sum of the four inputs for each gene, and then applied the same operation with a sigmoid activation function to compute the attention scores π_1_. The attention scores π_1_ is introduced here to let BioXNet learn the contributions of each input visibly. The output of the first layer is the attention-transferred output plus the original output followed by the tanh activation. We also utilized the dropout and batch norm operations to overcome overfitting as shown in equation (1).

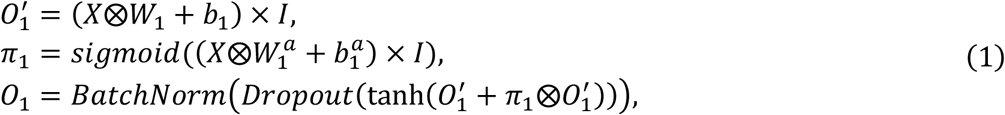

where *W*_1_ ∈ *R*^*n*×4^ and 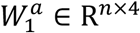 denote the weights; *b*_1_ ∈ *R*^*n*×4^ and 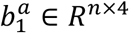 are the bias; and ⨂ denotes the element-wise multiplication; *I* ∈ *R*^4×1^ is a matrix with all elements equal to 1.

The following layer is restricted to the gene-pathway or pathway-pathway relationships using a mask matrix *M*. For layer *l*, suppose the number of input and output biological entities are 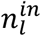 and 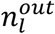, respectively. The mask matrix, denoted as 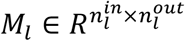, is a binary matrix where a value of 1 indicates the presence of a biological relationship from a node in the previous layer to a node in the next layer. We employed the element-wise product of *M*_*l*_ and the weight matrix *W*_*l*_ to zero-out the connections that are not present in the Reactome pathway dataset. For the attention module in layer *l*, we utilized a linear layer followed by a sigmoid activation function. The weight for the attention layer is 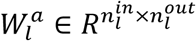, and the output is 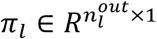 denoting the contribution scores of each biological entities in layer *l*. The final output of layer *l* is computed as follows:

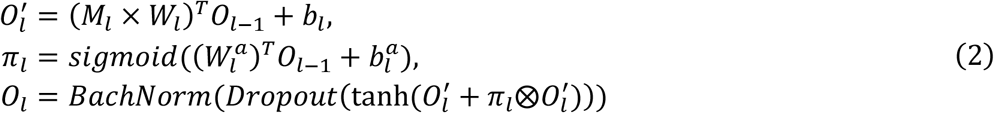

Follow the practice in P-Net^20^, BioXNet also utilize a prediction head layer for each hidden layer to allow each layer to be useful by itself. The prediction head layer is appended to the body of BioXNet and specifically designed for different tasks. The loss is a weighted combination of the loss from each prediction head layer, and it is computed as follows:

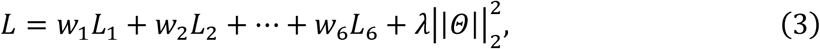

where *w*_*l*_ is a hyper-parameter representing the weight assigned to the loss of layer *l*. The last term in equation (3) is the *l*_2_ normalization to overcome overfitting, where *Θ* is the total parameters of BioXNet and *λ* is a hyper-parameter.

### 2. Training and weight transfer of BioXNet

BioXNet was initially trained on the cell line dataset, and the acquired knowledge is stored within the trained neural network. This knowledge can then be utilized to fine-tune drug response predictions on the patient dataset. The drug sensitivity on cell lines is measured by the half maximal inhibitory concentration (*IC*_50_), which indicates the effectiveness of the drug. The lower the *IC*_50_ value, the more effective the drug is considered to be. To meet the requirements for the model interpretation, we converted the *IC*_50_ values to *pIC*_50_, which is the negative logarithm of *IC*_50_. We utilized a linear layer with a single output as the prediction head layer. For layer *l*, the loss *L*_*l*_ is the root-mean-square error (RMSE) as shown in equation (4):

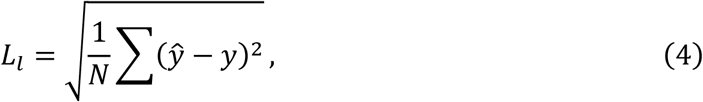

where *y* and 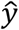 denotes the true and predicted *pIC*_50_, respectively.

After completing the training of BioXNet on the cell line dataset, we transferred the acquired knowledge to the clinical response prediction task for patients. This was achieved by initializing the weights of BioXNet using the model weights obtained from training on the cell line dataset. In this study, the clinical response prediction task involves a multi-class classification problem. The clinical responses are evaluated by Response Evaluation Criteria in Solid Tumors (RECIST)^29^ and categorized into four degrees: Complete Response (CR), Partial Response (PR), Stable Disease (SD), clinical Progressive Disease (PD). To represent these clinical responses numerically, the following values are assigned: CR is denoted as 3, PR as 2, SD as 1, and PD as 0. We utilized a linear layer with an output dimension of 4 and apply a SoftMax activation function as the prediction head layer. The loss here is the cross-entropy loss which is computed as follows:

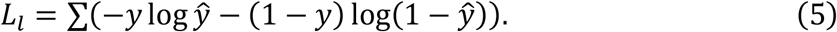

### 3. Datasets

#### Reactome

Reactome^25^ pathway database provides a hierarchy network of molecular transformations and reactions in the human body, including signal transduction, transport, metabolism and so on. We have curated a hierarchy network with 12,794 nodes including 9,375 genes and 3,419 pathways, which is connected by 36,685 biological relationships.

#### Genomics of Drug Sensitivity in Cancer (GDSC)

GDSC^26^ is the largest public dataset for information on drug sensitivity and bio-markers in cancer cell lines. The drug sensitivity in GDSC is measured by LN_IC50, the logarithm of *IC*_50_ values, which can be converted to *pIC*_50_ used in this study. After curation, we collected 135,353drug-cell line responses across 219 drugs and 479 cancer cell lines. The genomic profiles of cancer cell lines are available at Cancer Cell Line Encyclopedia (CCLE)^47^. We have collected the copy number variations (CNV), and the mutation of 14,812 genes. For the methylation data, to be consistent with that of TCGA, we collected methylation measured by 450K array from ^46^. Note that the original values in CNV are the normalized log2 ratios and we further converted these values to categorical copy number statuses following the same procedure^48^ in TCGA.

#### The Cancer Genome Atlas (TCGA)

TCGA^27^ is a research network that provided the genomic profiles of large numbers of human tumors to discover molecular aberrations at the DNA, RNA and so on. The clinical data in TCGA is relatively limited compared to the rich genomic profiles available. We have retained only the patients with available drug response, resulting in 2,765 drug-patient response data points across 1,265 patients and 112 drugs. Additionally, we have collected methylation data (measured by the 450K array), CNV information, and mutation information for the same 14,812 genes as in GDSC.

#### Drug-target

The drug-target information is binary, indicating whether a drug targets a specific gene. For drugs existed in GDSC or TCGA, we first utilized the DrugBank ID to identify their target genes. DrugBank^49^ is a comprehensive drug database including the chemical and biological knowledge of various drugs. For the remaining drugs without a corresponding DrugBank ID, we used their PubChem ID to find the target genes provided in our previous research^50^. In total, we collected 1,714 drug-target associations for 569 genes among the 121 drugs present in TCGA. In the case of GDSC, we curated 3,955 drug-target associations across 229 drugs and 1,059 genes. We categorized drugs into targeted therapies and chemotherapies based on the FDA classification (https://www.cancer.gov/about-cancer/treatment/types/targeted-therapies/approved-drug-list#targeted-therapy-approved-for-bladder-cancer).

#### Data preprocess

In the cell lines and patients from GDSC and TCGA, we limited our analysis to coding genes approved by the HUGO Gene Nomenclature Committee (HGNC)^51^.

We aggregated mutations at the gene level, focusing on nonsynonymous mutations in accordance with previous research on mutational significance^20^. Consequently, we excluded silent, intron, 3’ untranslated region (UTR), 5’ UTR, RNA, and long intergenic non-coding RNA (lincRNA) mutations from our analysis.

### 4. Experimental setup

#### Baselines

The baselines utilized in this study include a Dense neural network (Dense), P-Net, and ElasticNet. The Dense neural network refers to a fully connected neural network that has the same architecture as BioXNet. We also introduced P-Net^20^, a similar biological-inspired neural network used for cancer state prediction. Note that P-Net only considered genetic variations of genomic profiles (mutation and copy number variation). We modified P-Net by integrating drug-target information using the same method as BioXNet for drug response prediction. We fed Dense and ElasticNet the same input as BioXNet to have a fair comparison.

#### Evaluation metrics

The drug sensitivity prediction on cell lines is a regression task and the performance of BioXNet and baselines on this task is evaluated using RMSE and Pearson correlation. For the drug response prediction task on RECIST on TCGA, we used macro-averaged F1-score.

#### Implementation

BioXNet and baselines are implemented using Python 3.11. We employed a five-fold cross-validation with an 8:1:1 splitting ratio for training, validation, and testing. Performance results were reported based on the testing datasets. The source codes and data of BioXNet is available at https://github.com/JasonJYang/BioXNet.

### 5. Interpretation

#### DeepLIFT

DeepLIFT^28^ is a widely used attribution method for generating a sample-level contribution score for each feature. This method backpropagates the contributions of all neurons in the network to every input feature. The contribution is defined as the difference between the ‘reference activation’ *t*_0_ to the actual output *t*. Given a set of nodes 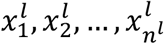 in one layer *l* in BioXNet, DeepLIFT computes the difference Δ*t* = *t* − *t*_0_ as 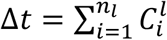, where 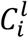 is the contribution score of node 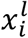 in the layer *l* given a specific input. In this study, the ‘reference activation’ *t*_0_ for the drug sensitivity prediction task on the cell line dataset (*pIC*_50_ prediction) is defined as value 0. For the drug response prediction task on the patient dataset, *t*_0_ is defined as ‘Complete Response’. Therefore, a positive contribution score 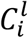 indicates a larger feature corresponds to a higher *pIC*_50_ value or a higher possibility to get ‘Complete Response’. This implies that the drug is more effective for the cell line or patient. Following the procedure in P-Net^20^, we aggregated the sample-level contribution scores by taking the absolute value of the summed contribution scores over specific samples and normalized the contribution scores using the node degrees. We implemented DeepLIFT to BioXNet using the Captum^52^ package.

#### Interpretable paths

Given an input drug-sample pair, BioXNet is capable of generating an interpretation graph (referred to as *G*_*I*_) that shares the same structure as the Reactome network used for training. In *G*_*I*_, nodes are assigned contribution scores obtained from DeepLIFT, while edges are assigned model training weights. To identify paths that best represent the interpretations for the drug response to a specific sample, we have proposed a random walk-based method considering both the node contribution scores and edge weights. Specifically, we created two virtual nodes (‘sample’ and ‘outcome’) into *G*_*I*_ as the starting and ending node, respectively. Starting from the ‘sample’ node, we randomly selected its neighbors as the next node considering each neighbor’s probability. This process was repeated until we reached the ‘outcome’ node, resulting in the desired path. Given one node *x*_*i*_ and one of its neighbor 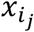, the probability is computed as the absolute value of the product of the node contribution score 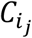 and the link weight 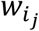 learned in BioXNet, that is 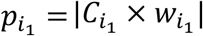. We repeated this process 10,000 times and recorded the frequency of each path. The interpretable paths are defined as the top 1% paths considering their frequency. This approach allows us to prioritize the most frequently occurring paths, providing a reliable and representative set of interpretable paths.

#### Readable interpretations

To further enhance the readability of the interpretations of BioXNet, we applied ChatGPT to generate readable interpretations based on the identified interpretable paths. Specifically, we utilized GPT-3.5 Turbo model with temperature 0 to ensure deterministic responses. The specific prompt used for this purpose is “*You are a professional biologist, especially at the field of drug response and mechanism of drug action. I have some paths generated from a model to indicate how the genomic profiles and drug targets affect the drug response to some cancer. Some paths starting from a drug, through several genes, biological pathways. And some paths starting from some modification (including methylation, copy number variation, and mutation) of genomics of some cancer patients, through genes, biological pathways. Now you need to combine these paths together to generate readable explanations to show how the genomic profiles and drug targets affect the drug response to this cancer. You only need to generate readable sentence and cannot change the meaning of these terms. Here are the paths:*” We developed an online service for users to investigate the interpretations generated by BioXNet, which is available at https://huggingface.co/spaces/Jayet010/bioxnet.

### 6. Statistical analysis

In the *pIC*_50_ prediction task (Figure 2a and 3f), *p*-values are determined using a two-sided test in the Pearson correlation. To ascertain *p*-values for performance and importance scores across various tissues or input data types (Figure 2d and Figure 3a-f, we employed a t-test. For the micro-F1 scores on individual tissues (Figure 2e), we implemented a bootstrapping statistical test with 10,000 samples, and the significance of the difference in score medians was tested.

## Acknowledgements

This work was supported by the Research Grants Council of the Hong Kong Special Administrative Region, China (Grant Nos. 11218221, C7154-20GF, C7151-20GF and C1143-20GF).

## Notes

https://github.com/JasonJYang/BioXNet

### Competing Interest Statement

The authors have declared no competing interest.

