## Supplementary Information for "BioXNet: a biologically inspired neural network for deciphering anti-cancer drug response in precision medicine"

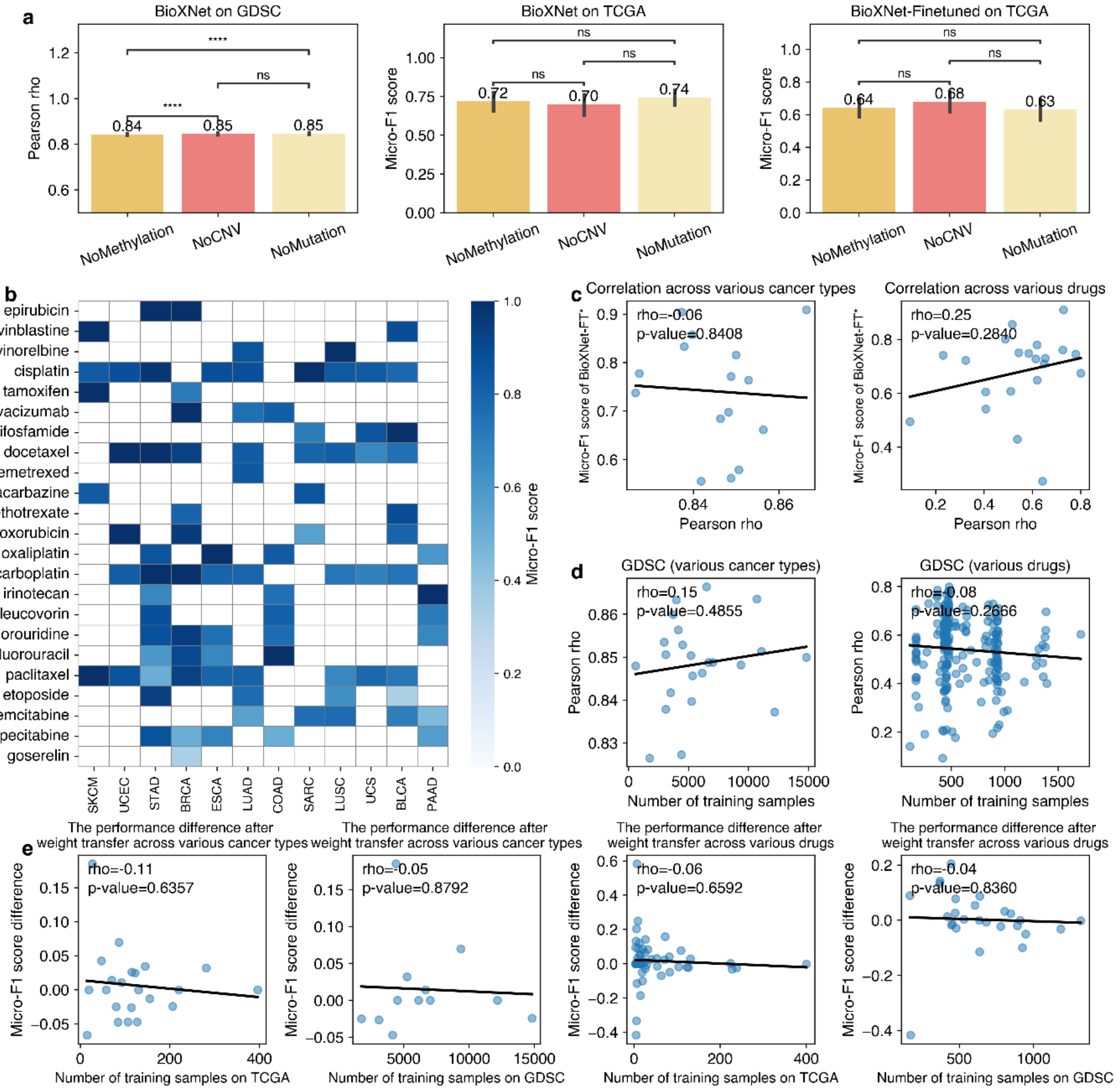


**Figure S1.** Evaluating the Performance of BioXNet. **a.** Comparison of BioXNet's performance using various input types. **b.** Heatmap displaying the performance of BioXNet-Finetuned in a clinical setting, considering different drug-cancer pairs involving Food and Drug Administration (FDA)-approved drugs. BioXNet-Finetuned represents the model that adopts weights transferred from BioXNet trained in a preclinical setting. Empty cells in the heatmap indicate non-existent drug-cancer pairs. **c.** Correlation analysis of BioXNet's performance in preclinical (measured by Pearson correlation rho, x-axis) and clinical settings (measured by micro-F1 score, y-axis) across diverse cancer types (left) and drugs (right). Note that the performance in preclinical settings was derived from BioXNet-FT* (BioXNet-Finetuned). **d.** Correlation analysis between BioXNet's performance and the number of training samples across various cancer types (left) and drugs (right). **e.** Investigation of the correlation between the improvement in performance in clinical settings (measured by micro-F1 score) after weight transfer and the number of training samples in The Cancer Genome Atlas (TCGA) or Genomics of Drug Sensitivity in Cancer (GDSC). The $p$-values in **a** were derived from t-test with annotation legends: $ns$: $p > 0.05$; $*$: $0.01 < p \leq 0.05$; $**$: $0.001 < p \leq 0.01$; $***$: $0.0001 < p \leq 0.001$; $****$: $p \leq 0.0001$.

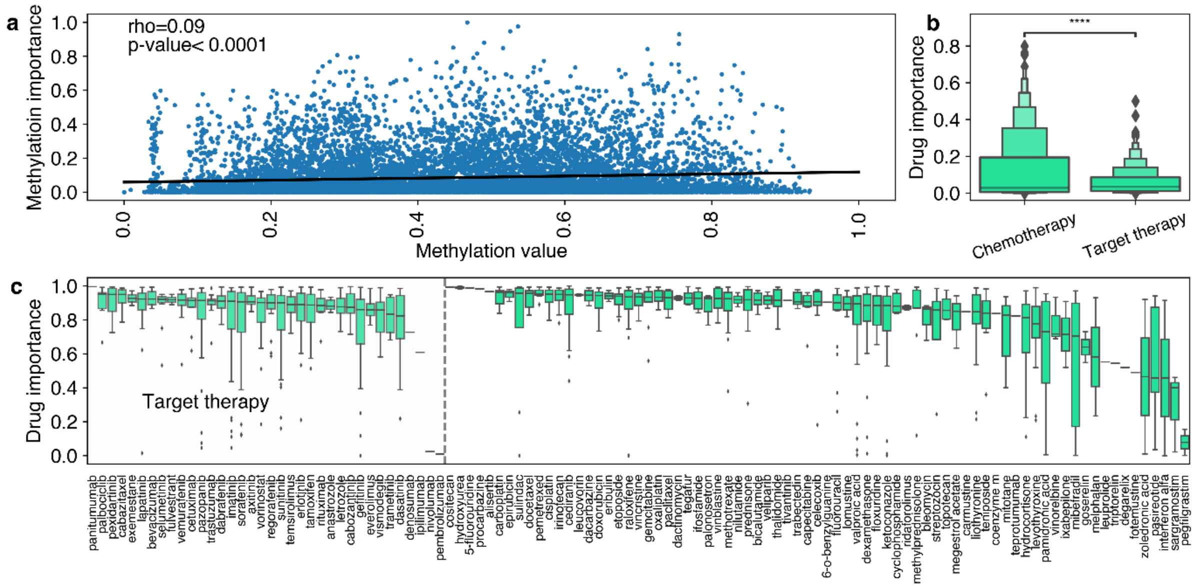


**Figure S2.** Investigating the Interpretability of BioXNet in Clinical Settings. **a.** Correlation analysis between methylation values and corresponding methylation importance scores across all cancer types for all genes. **b.** Comparison of drug importance scores for targeted therapy drugs and chemotherapy drugs in clinical settings. **c.** Analysis of drug importance scores in clinical settings across various drugs. The $p$-values in **c** were derived from t-test with annotation legends: $ns$: $p > 0.05$; $*$: $0.01 < p \leq 0.05$; $**$: $0.001 < p \leq 0.01$; $***$: $0.0001 < p \leq 0.001$; $****$: $p \leq 0.0001$.
